# Systems-level analysis of peripheral blood gene expression in dementia patients reveals an innate immune response shared across multiple disorders

**DOI:** 10.1101/2019.12.13.875112

**Authors:** D Nachun, EM Ramos, A Karydas, D Dokuru, F Gao, Z Yang, V Van Berlo, R Sears, J Kramer, AL Boxer, H Rosen, BL Miller, G. Coppola

## Abstract

The role of peripheral inflammation in dementia is an important but complex topic. We present here the largest cohort of peripheral blood gene expression data ever assembled from patients with dementia and matching controls. Importantly, this cohort includes individuals from a diverse set of dementia disorders, including Alzheimer’s Disease (AD), mild cognitive impairment (MCI), and multiple disorders within the frontotemporal dementia (FTD) spectrum. We found strong transcriptional evidence of an innate immune inflammatory response, mediated by monocytes and neutrophils, in AD, MCI, and two FTD subtypes, PSP and nfvPPA. This transcriptional inflammatory response is enriched for genetic risk for AD, in part because it is also enriched for microglial genes, which have previously been implicated in AD risk. Finally, we show that this transcriptional response is strongly enriched for binding of the transcription factors PU.1 and RELA, which have previously been linked to AD risk and progression.

## Introduction

Neurodegenerative forms of dementia constitute a heterogeneous group of diseases characterized by progressive loss of cognitive function driven by dysfunction and loss of neurons and glia in the central nervous system (CNS). Clinicians categorize dementias by the type of cognitive dysfunction observed in patients, such as memory loss, speech difficulties, or decline in executive function [1]. Pathological studies have identified specific proteinopathies such as accumulation of amyloid beta, tau, TDP-43, FUS, and α-synuclein in certain brain regions which have some association with clinical symptoms [2, 3]. Genetic studies have also implicated both rare and common variants in risk for dementia, some of which may be linked to associated proteinopathies. Notably, several genes harboring disease-associated variants, such as TREM2 [4], GRN [2], and others [5] are not directly associated with proteinopathy, but rather are involved in inflammatory response and innate immunity and are primarily expressed in microglia, implicating inflammation in the CNS as relevant to dementia pathophysiology [6–8].

Comparatively little is known about whether peripheral inflammation, in particular inflammation mediated by white blood cells (WBCs), is altered in dementia patients. Previous studies, often including small numbers of samples, have reported some evidence of increased peripheral inflammation in the most common form of dementia, Alzheimer’s disease (AD) [9], as well as in the most common neurodegenerative motor disorder, Parkinson’s disease [10], while it is unknown whether peripheral inflammation is altered in the spectrum of disorders associated with frontotemporal dementia (FTD), including behavioral variant FTD (bvFTD), semantic variant and nonfluent variant primary progressive aphasia (svPPA and nfvPPA), progressive supranuclear palsy (PSP), and corticobasal syndrome (CBS). We studied the largest cohort of peripheral blood gene expression data ever assembled from patients with dementia and matching controls, in order to determine whether a transcriptional response associated with disease status was detectable in RNA from peripheral blood.

## Results

Peripheral blood was collected from 1387 individuals for gene expression analysis. Gene expression was quantified using Illumina HT12 v4 microarrays. We only considered subjects with diagnoses of control, AD, MCI, bvFTD, svPPA, nfvPPA, PSP or CBS, and also removed RNA samples with RIN < 6.0. After filtering by these criteria, we had 981 RNA samples (281 control, 269 AD, 172 MCI, 82 bvFTD, 52 PSP, 46 nfvPPA, 44 svPPA, 35 CBS, Table 1).

**Table 1.** Demographic summary of subject data.

| Diagnosis | Mean Age | SD Age | Female | Male | Total |
| --- | --- | --- | --- | --- | --- |
| Control | 71.5 | 10.1 | 156 | 125 | 281 |
| AD | 68.8 | 9.8 | 144 | 125 | 269 |
| MCI | 69.4 | 10.9 | 78 | 94 | 172 |
| bvFTD | 61.2 | 9 | 37 | 45 | 82 |
| PSP | 70.2 | 6.6 | 27 | 25 | 52 |
| nfvPPA | 67.7 | 7.6 | 27 | 19 | 46 |
| svPPA | 65.7 | 7.2 | 21 | 23 | 44 |
| CBS | 67 | 7.6 | 20 | 15 | 35 |
| **Total or Average** | **67.7** | **8.6** | **510** | **471** | **981** |

### Differential Expression

#### AD and MCI

We used Bayesian linear modeling (see Methods) to identify transcripts which were differentially expressed (DE) between controls and patients with AD and MCI. We identified a significant confound between age and diagnosis in the full dataset which we resolved by stratifying our samples so that only subjects between the ages of 60 and 90 were included. After this stratification process, as well as the removal of expression outliers, we included 229 control samples, 198 AD samples, and 124 MCI samples in our analysis. At a significance cutoff of logBF > 0.5 and posterior probability > 0.95, we identified 444 DE genes between AD and controls (global FDR = 0.0073, Figure 1a, Table S1), 451 between MCI and controls (global FDR = 0.0057), and 280 between AD and MCI (global FDR = 0.012). Enrichment analysis of differentially expressed genes (Figure 1b) revealed a significant and highly specific enrichment for neutrophil degranulation (GO:0043312), a key component of the innate immune response, in upregulated genes in AD vs. Control (39 genes, logBF = 11.20) and MCI vs. Control (28 genes, logBF = 4.31), and a weaker enrichment in AD vs. MCI (13 genes, logBF = 1.91).

**Figure 1.**
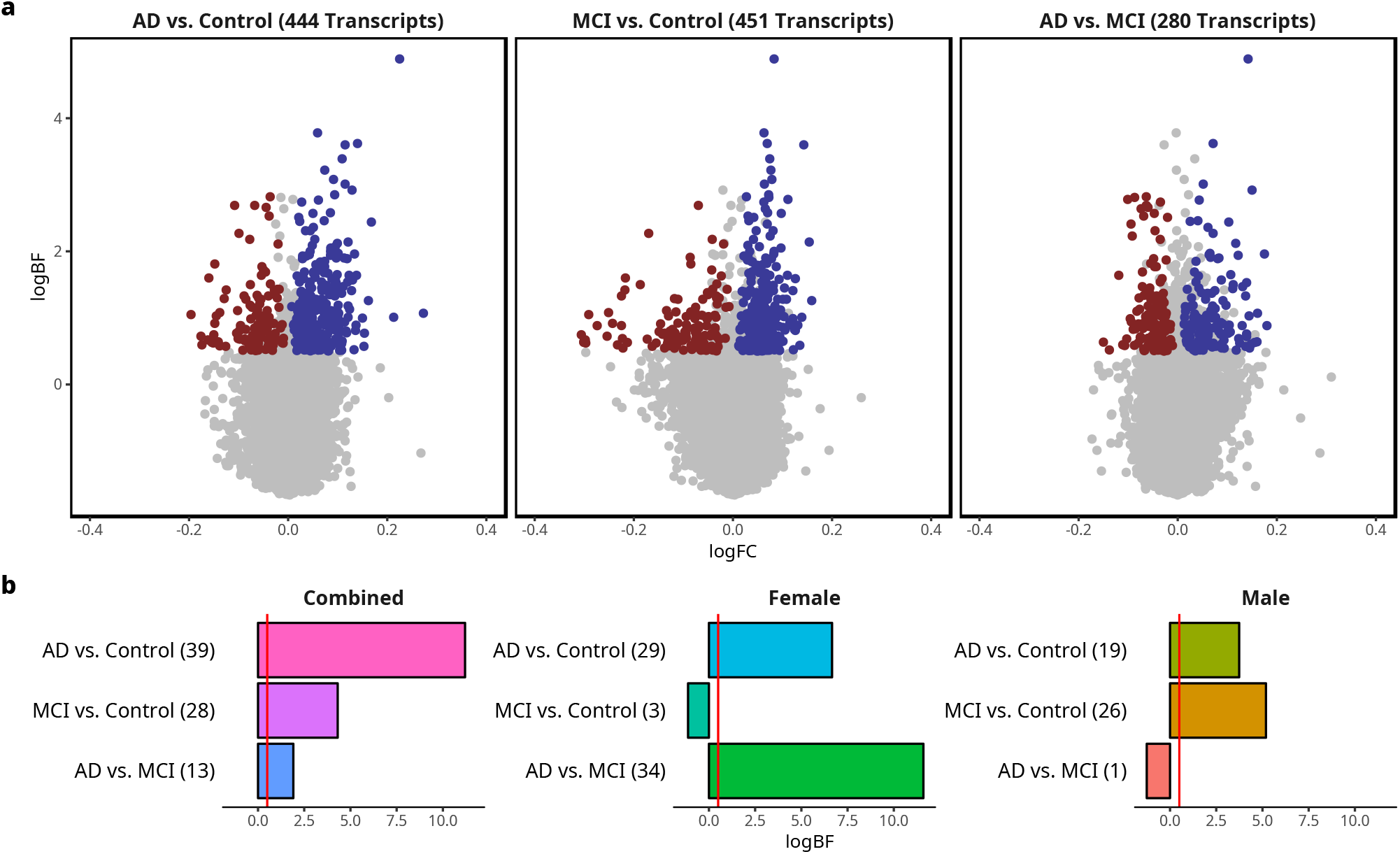
Differential expression analysis on AD, MCI and control subjects identifies an innate immune signature in blood. **a** Volcano plots of the log fold change (logFC) in gene expression on the x-axis versus the log10 Bayes Factor (logBF) on the y-axis, with downregulated genes in blue and upregulated genes in red. Only genes with a significant pairwise probability greater than 0.95 were colored. The contrast and number of DE genes are shown in the plot titles. **b** Bar plots of enrichment of significantly upregulated genes for neutrophil degranulation (GO:0043312) with the logBF on the x-axis.

#### Sex differences in AD and MCI

Studies of the epidemiology of dementia, particularly AD, have identified sex differences in disease risk including consistent evidence that females have a higher risk of developing AD [11]. We partitioned our samples by sex to determine if there were differences in the inflammatory response we observed in the full dataset between males and females. Each dataset was preprocessed separately, resulting in 251 male samples (94 controls, 93 AD, 64 MCI) and 269 female samples (123 controls, 94 AD, 52 MCI). The number of DE genes in both females (AD vs. Control: 355 genes, global FDR = 0.0076; MCI vs. Control: 293 genes, global FDR = 0.011; AD vs. MCI: 468 genes, global FDR = 0.0055, Figure S1a, Table S1) and males (AD vs. Control: 280 genes, global FDR = 0.0076; MCI vs. Control: 393 genes, global FDR = 0.011; AD vs. MCI: 157 genes, global FDR = 0.0055, Figure S1b, Table S1) was comparable to the analysis on the combined dataset. However, the enrichment in neutrophil degranulation (Figure 1b) in upregulated genes in MCI vs. Control was observed in males (26 genes, logBF = 5.19) but not in females MCI vs. Control (3 genes, logBF = −1.13). This indicates that, while the MCI vs. control comparison in males and females shows a similar numbers of DE genes, male MCI subjects exhibit an inflammatory response in peripheral blood similar to that in AD subjects, whereas a comparable inflammatory response is not detectable in female MCI subjects.

#### FTD disorders

We applied the same linear modeling approach to identify transcripts that were differentially expressed between FTD disorders and control (268 controls, 75 bvFTD, 53 PSP, 45 nfvPPA, 43 svPPA, 35 CBS). A significant age confound was also observed with diagnosis in this subset of the data which could not be resolved using stratification and was instead removed using residualization (see Methods). Using the significance cutoffs previously described, we identified 175 DE genes in bvFTD vs. control (global FDR = 0.013, Figure S2a, Table S1), 189 genes in nfvPPA vs. control (global FDR = 0.016), 81 genes in svPPA vs. control (global FDR = 0.02), 257 genes in PSP vs. control (global FDR = 0.0073, Figure S2b) and 48 genes in CBS vs. control (global FDR = 0.023). Enrichment analysis of upregulated genes in each disease vs. control only revealed significant enrichment for neutrophil degranulation in PSP vs. control (18 genes, logBF = 2.08), and no significant enrichments in other diseases. We did not attempt to partition our FTD disorder data by sex because our samples sizes for these disorders were much smaller than for AD and MCI.

We next asked how similar the transcriptomic effects of each FTD-spectrum disorder were to the other disorders. To visualize this, we correlated the logFC values of disease vs. control for all genes in each FTD disorder with the corresponding logFC values in each of the other disorders and clustered the diseases based on these correlation values (Figure S2c). We found that nfvPPA and PSP, both known to be tauopathies [12, 13] clustered with each other and away from the other categories, suggesting the presence of a tau-related signal.

### Network Analysis

Our differential expression results showed that there was an enrichment for an innate immune transcriptional response, and also pointed to a possible effect of sex on MCI vs. control gene expression signatures. We used weighted gene co-expression network analysis (WGCNA) to identify clusters of co-expressed transcripts (also known as modules) which are often highly enriched for specific biological pathways [14–20]. The gene expression pattern in a module across samples can be summarized by the first principal component of the expression values of all the genes in a given module, or eigengene. These eigengenes were analyzed using the same linear modeling approach used for differential expression, with the same significance cutoffs. We used this method on the same data subsets analyzed with differential expression to better understand the innate immune response signature we identified throughout our analyses.

#### AD and MCI

WGCNA identified 18 modules of co-expressed genes in the full dataset of AD, MCI, and control subjects. Disease status was a significant predictor for the magenta (logBF = 1.28, Figure S3a, Table S2) and brown (logBF = 0.723, Figure 2a) modules. Pairwise comparisons for the magenta module showed a significant increase in AD vs. control (diff. = 0.0085, pp. = 0.982) and MCI vs. control (diff. = 0.017, pp. = 1.0) and a marginally significant decrease in AD vs. MCI (diff. = −0.0081, pp. = 0.958). In the brown module (Figure 2a), AD (diff. = 0.012, pp = 0.998) and MCI (diff. = 0.011, pp = 0.991) were significantly increased vs. Control, while no significant difference was observed between AD and MCI (diff. = 0.0012, pp = 0.595). Both modules showed significant enrichment for neutrophil degranulation (magenta: 45 genes, logBF = 0.67, Figure S3e; brown: 123 genes, logBF = 25.2, Figure 2d) but clearly showed different expression patterns with regards to MCI.

**Figure 2.**
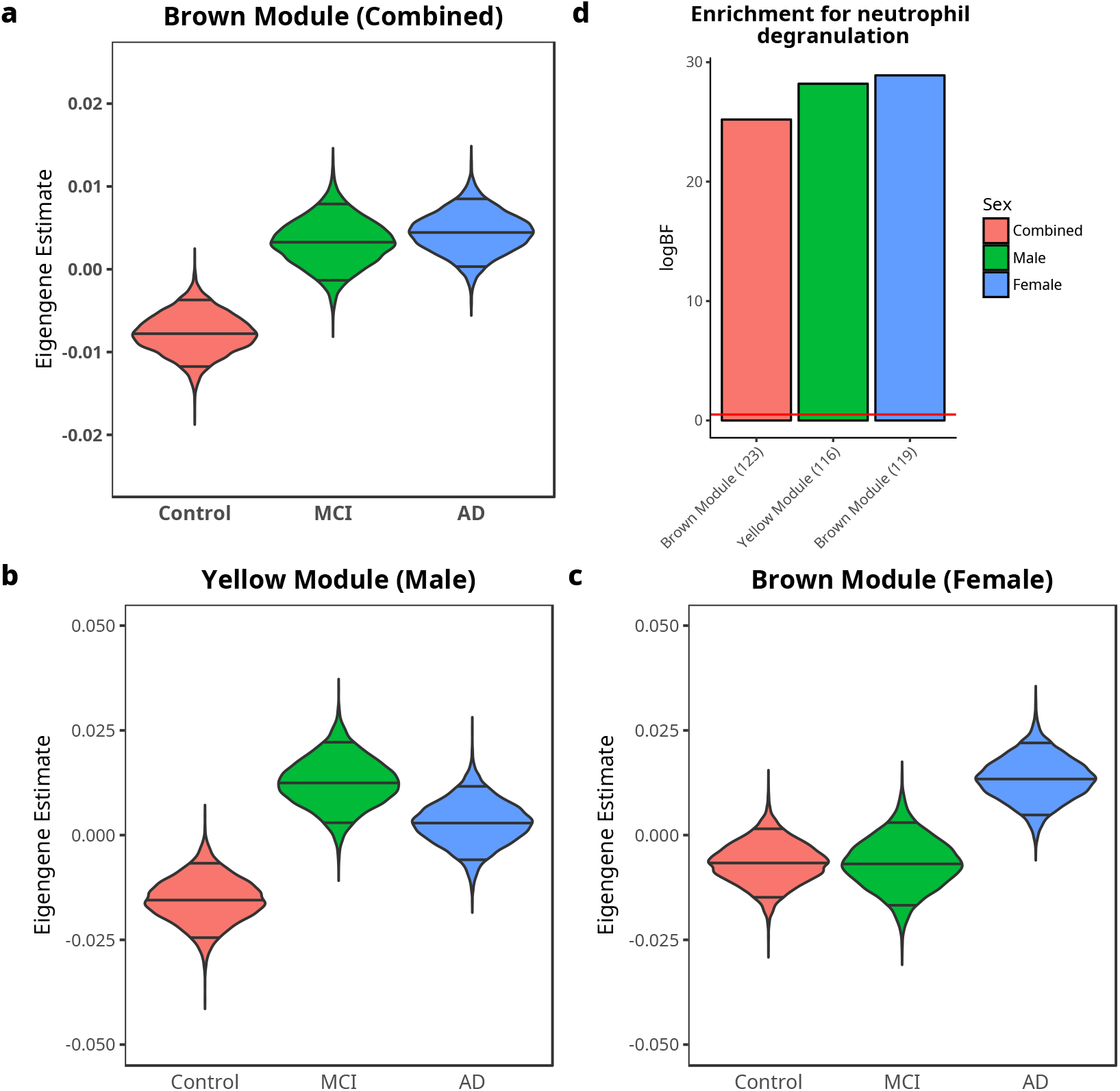
WGCNA identifies modules significantly associated with disease in AD, MCI and control subjects. **a-c** Violin plots of posterior estimate of mean eigengene values for each diagnosis, with the median and 5% and 95% quantiles indicated by lines. **c** Bar plot of enrichment of genes in each module for neutrophil degranulation (GO:0043312) with the logBF on the x-axis.

Since our previous analyses pointed to a possible effect of sex in expression patterns, we regenerated our co-expression network in males and females separately. In the male samples, we identified two out of 19 modules where diagnosis had a significant effect: the green module (logBF = 0.945, Figure S3b, Table S2) and the yellow module (logBF = 0.619, Figure 2b). The pairwise comparisons for the green module showed that it was significantly increased in both AD (diff. = 0.0159, pp. = 0.962) and MCI vs. control (diff. = 0.0312, pp = 1.0), but not significantly different for AD vs. MCI (diff. = −0.0155, pp. = 0.947). Similarly, the yellow module was also significantly increased in both AD (diff. = 0.0184, pp. = 0.980) and MCI vs. control (diff. = 0.0279, pp. = 0.998) but was not different between AD and MCI (diff. = −0.00965, pp. = 0.844). Enrichment analysis of both modules revealed that the yellow module was strongly enriched for neutrophil degranulation (116 genes, logBF = 28.2, Figure 2d) while the green module was only weakly enriched for the same pathway (51 genes, logBF = 0.901, Figure S3e).

In female samples we also identified two modules (out of 23) where diagnosis had a significant effect: the black (logBF = 1.69, Figure S3d, Table S2) and the darkgrey module (logBF = 1.57, Figure S3c). The pairwise comparisons for the black module found MCI vs. control was significantly increased (diff. = 0.0326, pp = 1.0) and AD vs. MCI was significantly decreased (diff. = 0.0345, pp = 1.0), but there was no significant difference between AD and control (diff. = −0.00198, pp = 0.60). In contrast, the darkgrey module was significantly increased in AD vs. control (diff. = 0.0185, pp = 0.988) and AD vs. MCI (diff = 0.0361, pp = 1.0) and was significantly decreased in MCI vs. control (diff. = −0.0178, pp = 0.967). We also found that the brown module, while not significant for model comparison (logBF = 0.212, Figure 2c), still showed a significant increase in AD vs. control (diff. = 0.020, pp. = 0.993) and AD vs. MCI (diff. = 0.020, pp. = 0.977) but no difference in MCI vs. control (diff. = −0.0001, pp. = 0.505). Enrichment analysis of these modules revealed significant enrichment in the darkgrey module (26 genes, logBF = 7.11, Figure S3e) and brown module (119 genes, logBF = 28.9, Figure 2d) for neutrophil degranulation but no enrichment for the same pathway in the black module (7 genes, logBF = −0.735, Figure S3e). These results confirmed our observation in the differential expression analysis that while both male and female AD subjects show evidence of increased inflammatory response, an increased inflammatory response in MCI is only observed in males.

#### FTD disorders

WGCNA in the cohort of FTD disorders and controls identified 14 modules, none of which identified disease status as a significant overall predictor. However, pairwise comparisons to controls identified the brown module as associated with PSP (upregulated, diff. = 0.0127, pp. = 0.982, Figure 3a, Table S2) and nfvPPA (upregulated diff. = 0.0107, pp. = 0.953). Enrichment analysis of this module found that it was strongly enriched for neutrophil degranulation (brown: 153 genes, logBF = 40.5). These results support the notion that a transcriptional signature associated with increased innate immune response is present in PSP vs. control but not in bvFTD vs. control, and provide some support for an increased innate immune response in nfvPPA vs. control, which could not be detected with differential expression.

**Figure 3.**
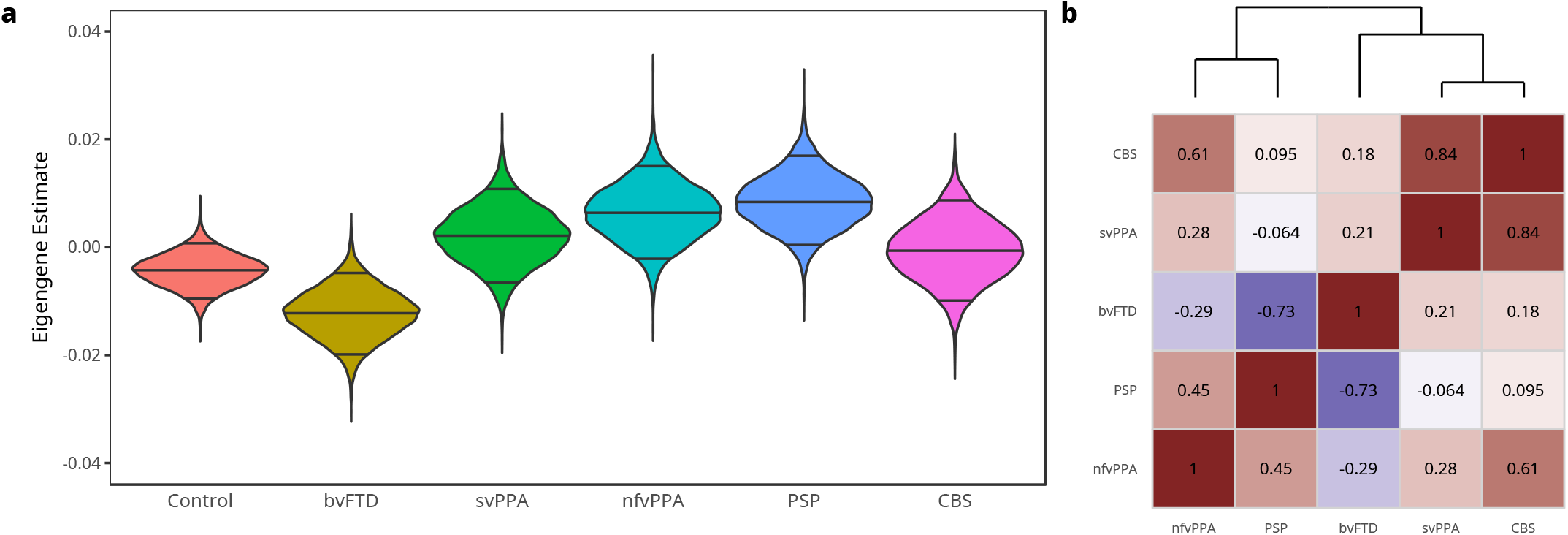
WGCNA identifies modules significantly associated with disease in FTD spectrum and control subjects. **a** Violin plot of posterior estimate of mean eigengene values for each diagnosis in the brown module, with the median and 5% and 95% quantiles indicated by lines. **b** Correlation plot of the pairwise correlation of the mean differences in eigengene values of FTD disorders vs. control, with the dendrogram on top showing hierarchical clustering of disorders.

Like we had previously done with the differential expression data in the FTD disorders, we correlated the mean differences between disease and control for each eigengene in one disease with the mean differences in the other diseases to understand how similar the disease effects were at the level of co-expression modules. We observed a disease clustering similar to that observed in the DE analysis: nfvPPA and PSP forming one cluster, svPPA and CBS forming another cluster, and bvFTD remaining unique (Figure 3b).

#### Enrichment of network modules for cell type specific genes in blood

Because we observed clear evidence of an innate immune response in all of our WGCNA networks, we wanted to determine whether our modules were enriched for genes specific to cell types in blood. We used the pSI [21, 22] tool to identify cell type-specific genes from an existing dataset [23] and then tested for significant overlap with the top 300 genes in each of our modules. As shown in Figure 4, the brown module in both the AD (neutrophils: 61 genes, logBF = 20.3; monocytes: 84 genes, logBF = 31.9) and FTD (neutrophils: 57 genes, logBF = 16.8; monocytes: 83 genes, logBF = 30) networks was clearly the most significantly enriched for cell type specific genes in neutrophils and monocytes. Importantly, in all analyses we identified other modules not affected by disease which are also enriched for neutrophils and monocytes, indicating that not all of the transcriptome associated with these cell types is affected by disease. We also show that the modules we found to be enriched for neutrophil and monocyte genes are not enriched for any other blood cell types (Table S3).

**Figure 4.**
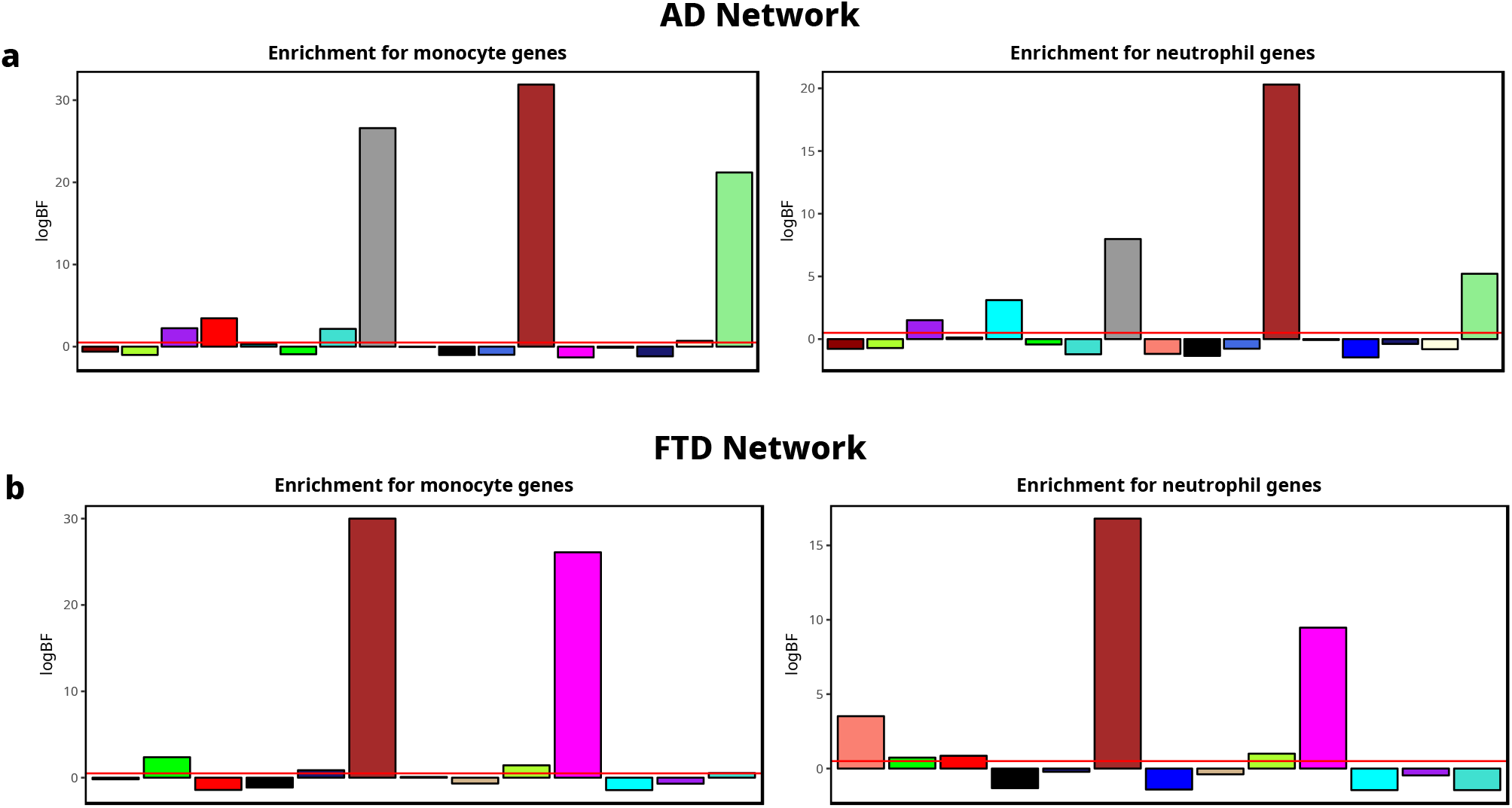
Disease-associated WGCNA modules are enriched for monocyte- and neutrophil-specific genes. **a,b** Bar plots of enrichment of the top 300 genes in each WGCNA module for monocyte- and neutrophil-specific genes in the AD and FTD networks. The logBF is on the y-axis and modules are ordered by the hierarchical clustering of their eigengenes. The number of genes in each overlap is in parentheses next to the module name in the legend.

### Cell Type Composition

One potential confounding factor in our analysis, that could have led to the identification of an innate immune response enrichment in upregulated genes in AD and FTD disorders, is a change in cell type composition. We used CIBERSORT [24] to estimate cell type composition and determine if there were any changes as a result of disease. This tool uses gene expression profiles from FACS-sorted cell types in peripheral blood to estimate the proportion of the most common types of cells seen in blood. Figure S4a shows the percent composition of cell types in the cohort of AD, MCI and control and Figure S4b shows the percent composition of cell types in the FTD disorders and controls. In the FTD disorders, the effect of age on cell type composition was removed with residualization before analysis. In both cohorts, no significant effect of diagnosis was seen on the proportions of any cell type, indicating that a change in cell type composition as determined by gene expression was not responsible for the innate immune response signal detected in the differential expression and network analysis.

### Enrichment for AD genetic risk

After identifying a consistent innate immune response signature in AD, MCI, and FTD disorders in our co-expression networks, we wanted to investigate the possibility that the immune response modules we identified were enriched for genes associated with the genetic risk for AD resulting from common genetic variation.

We used MAGMA [25], a tool designed to test gene sets for enrichment for genetic association with a trait using genome wide association study (GWAS) summary statistics. The GWAS we used was generated by the International Genomics of Alzheimer’s Project (IGAP) using 17,008 AD cases and 37,154 controls and genotyped or imputed 7,055,881 single nucleotide polymorphisms (SNPs) [26]. To completely remove any effects of ApoE, we removed all SNPs from the IGAP summary statistics within 1 centimorgan of rs7412.

We used the top 300 genes in each module from each of our networks as the gene sets to be tested for enrichment for genetic association. In the AD/MCI network, we found the brown (p < 0.004) module to be significantly and exclusively enriched for genetic risk for AD (Figure 5a). This trend holds in the FTD network (Figure 5b), where robust enrichment is exclusive of the brown module as well (p < 0.000121). We also tested whether this enrichment was specific to AD or was seen with other GWAS studies. We observed no enrichment, in either network, for the innate immune modules described above for genetic risk for amyotrophic lateral sclerosis [27], age of onset in Huntington’s Disease [28], type 2 diabetes [29], schizophrenia [30], or major depression [31], indicating that this enrichment for genetic risk is specific to AD (Table S4).

**Figure 5.**
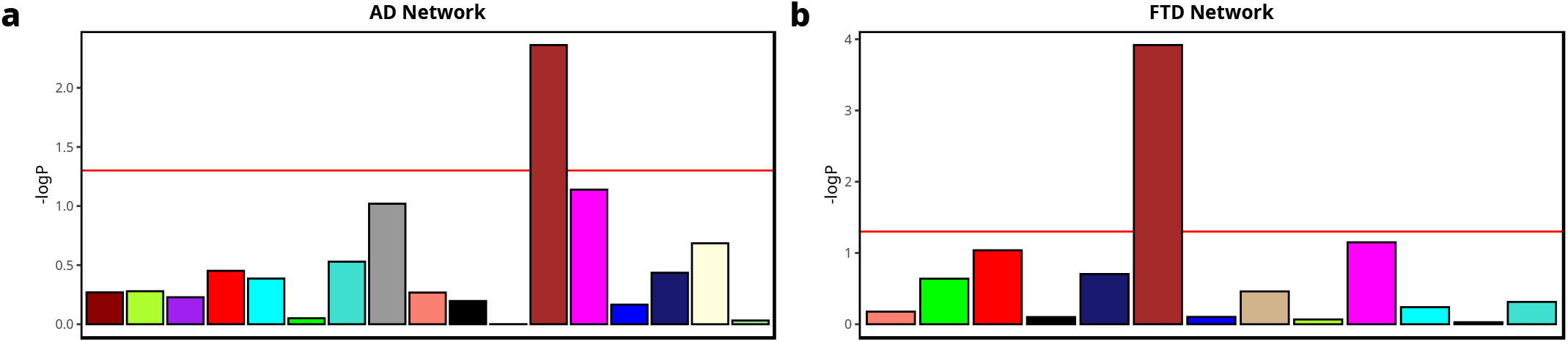
Disease-associated WGCNA modules are enriched for AD genetic risk. **a,b** Bar plots of enrichment of the top 300 genes in each WGCNA module for genetic risk for AD as estimated by MAGMA in the AD and FTD networks. The −log10 p-value (adjusted for multiple comparisons) is on the y-axis and modules are ordered by the hierarchical clustering of their eigengenes.

### Enrichment for microglia genes

After observing a significant enrichment for genetic risk in the modules associated with an innate immune response in our co-expression networks, we hypothesized that this could be because the immune response in the peripheral blood transcriptome overlaps with the transcriptome of microglia, which mediate innate immune responses in the CNS. To test this, we used a cell type-specific gene expression dataset collected from human brains [32] and the pSI tool [21, 22] to identify cell type-specific genes in microglia as well as neurons, astrocytes, oligodendrocytes, and endothelial cells and tested the same genes sets from network modules analyzed in MAGMA for enrichment for these cell type specific genes. In the both the AD network (26 genes, logBF = 5.92) and the FTD network (23 genes, logBF = 4.16), we found that the brown module was strongly enriched for microglia-expressed genes (Figure 6), and showed little or no enrichment for the other cell types in the CNS (Table S3). We note that in all cases other modules in the network also showed enrichment for microglia-specific genes, but these modules were not significantly increased in disease, suggesting that our analyses identify a subset of microglia-expressed genes which are associated with disease status, and detectable in peripheral blood.

**Figure 6.**
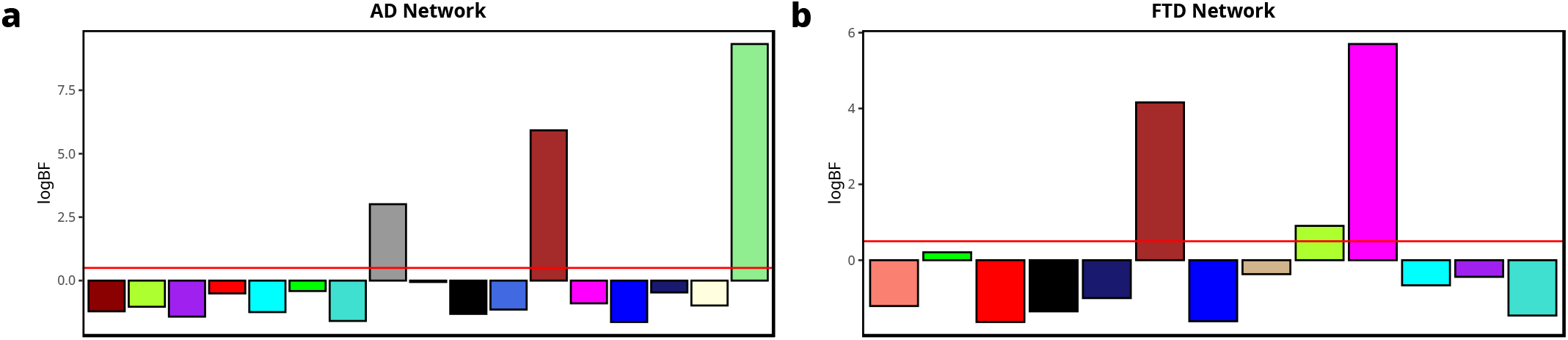
Disease-associated WGCNA modules are enriched for microglia-specific genes. **a,b** Bar plots of enrichment of the top 300 genes in each WGCNA microglial genes in the AD and FTD networks. The logBF is on the y-axis and modules are ordered by the hierarchical clustering of their eigengenes. The number of genes in each overlap is in parentheses next to the module name in the legend.

### Enrichment for transcription factor binding

Previous studies using co-expression networks have found that gene expression in some modules is regulated by transcription factors [16]. The simplest way of identifying enrichment of transcription factor targets in a co-expression module is an overlap analysis between the genes in the module and genes identified as targets of the transcription factor, like the approach used to identify enrichment for cell type specific genes. We used a more sophisticated approach that exploits the availability of ChIP-Seq data generated in myeloid leukemia cell lines (K562 and HL-60), which are biologically similar to monocytes and granulocytes. Similar to the approach used in MAGMA [25], we averaged log P-values of all ChIP-Seq peaks in the gene body and 1.5kb upstream of the transcription start site for each gene in our co-expression networks, and used linear models to compare the scores of genes in each of our modules to the rest of the genes in the network. We used ChIP-Seq data for 2 transcription factors known to be involved in myeloid lineage specification, SPI1/PU.1 and CEBPB, and 2 transcription factors important for inflammatory response in myeloid cells, RELA (a subunit of NF-κB) and JUN (a subunit of AP-1).

Figure 7 shows that in both the AD (logBF = 50.3) and FTD (logBF = 56.2) networks, the brown module was strongly enriched for binding of PU.1, and that the enrichment was also highly specific to that module. While enrichment for PU.1 binding is expected in myeloid cells given its importance in regulating myeloid lineage specification [33], this result is still intriguing given that a variant in PU.1 which reduces its expression was found to delay the onset of AD [34], and PU.1 gene expression is significantly increased in AD (logFC = 0.11, pp = 1.0), MCI (logFC = 0.083, pp = 0.98), and PSP (logFC = 0.18, pp = 1.0). Strong, highly specific enrichment in the brown module in both networks was also seen binding of the other myeloid lineage factor CEBPB (Figure S5), and like PU.1, its expression was significantly increased in AD (logFC = 0.06, pp = 1.0) and PSP (logFC = 0.1, pp = 0.98), but not MCI (logFC = −.006, pp = 0.6). RELA also showed very specific enrichment for the brown module while much less specific enrichment was seen for JUN, indicating that NF-κB but not AP-1 signaling is enriched in the brown module. A key adaptor protein for IL-1R and TLR signaling through NF-κB, MYD88, also showed significantly increased gene expression in AD (logFC = 0.057, pp = 0.99), MCI (logFC = 0.06, pp = 0.99), and PSP (logFC = 0.11, pp = 1.0).

**Figure 7.**
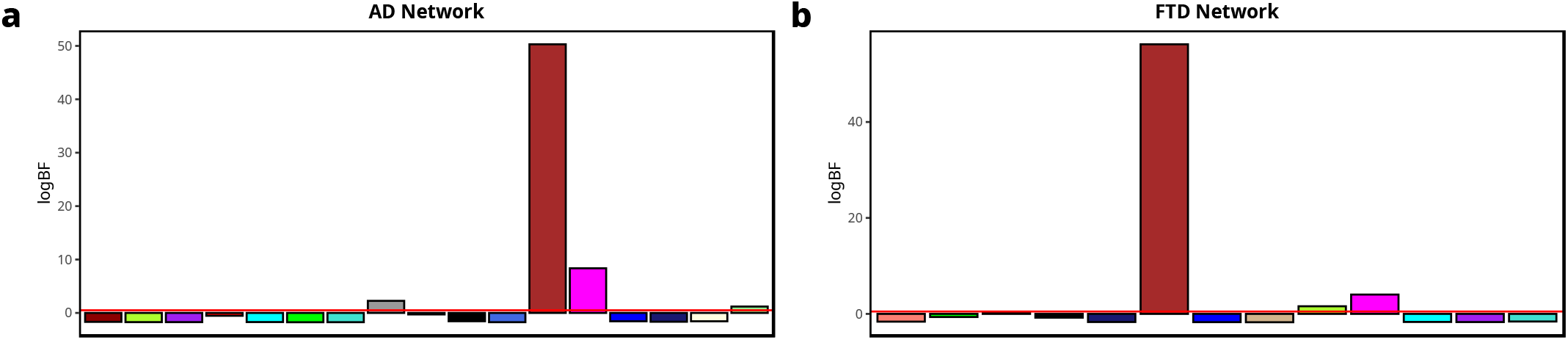
Disease-associated WGCNA modules are enriched for PU.1 binding. **a,b** Bar plots of enrichment of the top 500 genes in modules in the AD and FTD networks for PU.1 binding.

## Discussion

Our comprehensive analysis of peripheral blood gene expression in AD, MCI and FTD disorders detected a transcriptional signature indicative of an increased inflammatory response in monocytes and neutrophils in AD, MCI in males, and PSP, and modest evidence for this increase in nfvPPA. Whether this response is also increased in svPPA and CBS is unclear, although their transcriptional profiles are correlated, whereas bvFTD may show evidence of decreased inflammation. We further showed that the inflammatory response we identified using network analysis is significantly and specifically enriched for genetic risk for AD, even when considering the inflammatory response seen in nfvPPA and PSP. We showed that this genetic enrichment is likely driven by the strong overlap between the transcriptional inflammatory response we observe in peripheral innate immune cells with genes that are expressed in microglia. Finally, the genes in these innate immune modules were strongly enriched for binding of several transcription factors associated with innate immune inflammatory responses, particularly PU.1. We also demonstrated that this transcriptional response is not driven by differences in blood cell type composition between diseases and control.

The most likely reason for the correlation observed between transcriptional changes in innate immune cells in blood and microglia in the CNS is that the inflammatory signals responsible for activating the microglial response are also present in blood. Inflammatory peptides are increased in the serum of AD patients [9], and there is evidence that monocytes are responsible for clearing circulating amyloid beta [35], tau [36], alpha-synuclein [37], and TDP-43 [38]. We saw no clear evidence of enrichment of neuron-, astrocyte- or oligodendrocyte-specific genes in any of the co-expression modules we identified in blood, which is unsurprising given that these other CNS cell types do not have analogous cell types in blood. However, this is consistent with post-mortem gene and protein co-expression networks in AD brains [39], which showed no evidence of enrichment for genetic AD risk in genes specific to neurons and astrocytes, and only weak enrichment in gene specific for oligodendrocytes. By contrast, many psychiatric disorders show strong enrichment for genetic risk in genes specific to neurons [40] and no enrichment in microglia, which may limit the value of studying peripheral blood gene expression in those disorders.

The observation that a transcriptional inflammatory response in PSP and nfvPPA would be enriched for AD genetic risk is intriguing because it implies that there is some overlap in inflammatory signaling across diseases. Currently available GWAS studies for FTD disorders are very underpowered, but these results predict that, at least for PSP and nfvPPA, some genetic risk may be mediated through microglial genes, similar to what has been reported in AD. Similarly, the fact that bvFTD does not show an increased inflammatory response in blood and the lack of enrichment for ALS genetic risk in any of our co-expression modules, a disorder that is comorbid with bvFTD and shares many genetic risk factors [2], suggests that microglial inflammatory responses are not as relevant to bvFTD pathology as they are in AD.

The sex difference we identified in the inflammatory response in MCI patients may be related to the sex differences in the clinical presentation of MCI. Previous studies have found that females show less memory impairment than males with an equivalent level of neurodegeneration [41, 42], which may lead to underdiagnosis of MCI in females. In our data, we saw no evidence of an inflammatory response in female MCI subjects, but saw that in male MCI subjects, the inflammatory response was as strong as that seen in AD subjects. This could be due to different prodromal AD trajectories in females whereby impaired cognition would be detectable only when it has progressed to AD itself, while in males it would be detectable during earlier stages as MCI, before progression to AD.

One of the most important questions that emerges from these findings is whether the innate inflammatory response is pathological or protective. This is one of the central topics of debate in neurodegeneration research, with contradictory evidence showing that inflammation in the CNS worsens the progression of neurodegeneration, and that it slows it. The role of peripheral inflammation is even less clear. However, the observation of increased PU.1 expression in the peripheral transcriptomic inflammatory response does provide some insight into this question. A common haplotype in this gene was found to reduce expression of PU.1 and was found, at an epidemiological level, to delay the onset of AD [34]. PU.1 is known to regulate numerous AD-related genes in microglia [43], including TREM2, ABCA7 and CD33, and to regulate the differentiation of macrophages from monocytes [44]. Taken together, these data imply that the PU.1-mediated transcriptional response we observe may ultimately lead to increased macrophage differentiation and infiltration into the CNS in response to inflammatory signaling, and that inhibiting this process may be protective against disease progression. There may be a similar effect from increased expression of CEBPB, and the additional evidence of NF-κB signaling, which also regulates many AD risk genes [45] and can be activated by IL-1B signalling [46], further reinforces this idea. Such a model has been proposed before, based primarily upon evidence obtained from mouse models of AD [35, 47] with small sample sizes. This study provides support for this model using a large sample taken directly from human patients with neurodegenerative disease.

## Methods

### Subjects

Table 1 summarizes subject demographics. All subjects provided consent for using their data for research purposes, and the data collection protocols were approved by the institutional review boards of UCLA and UCSF.

### Sample preparation

Peripheral blood was collected in Paxgene tubes and frozen before RNA extraction, which was performed using a semi- automated system (Qiacube). Subjects were not specifically instructed to fast or refrain from exercise, and the time of collection was not uniform. RNA quantity was assessed with Nanodrop (Nanodrop Technologies) and quality with the Agilent Bioanalyzer (Agilent Technologies), which generated an RNA integrity number (RIN) for each sample. Total RNA (200 ng) was amplified, biotinylated and hybridized on Illumina HT12 v4 microarrays, as per manufacturer’s protocol, at the UCLA Neuroscience Genomics Core. Slides were scanned using an Illumina BeadStation and signal extracted using the Illumina BeadStudio software (Illumina, San Diego, CA).

### Quantification and statistical analyses

#### Gene expression array preprocessing

Illumina HT-12 v4 microarrays were preprocessed using the lumi pipeline [48] as previously described [49]. Expression values were normalized using the variance-stabilized transformation [50] and robust spline normalization was used for inter-array normalization. Probes with a detection score p-value greater than 0.01 were dropped, as were probes that were unannotated. Duplicated probes for the same transcript were dropped using the collapseRows function [51] from the WGCNA package. Outliers were removed based on connectivity z-scores [52]. Batch correction was performed using ComBaT [53] and any batch with less than 8 samples was dropped to allow for more robust estimation of batch effects. Subsets of data such as those partitioned by sex were preprocessed separately, meaning that different sets of samples were identified as outliers, and different batches were dropped based on the number of samples per batch in a given subset of data.

#### Linear Modeling and Residualization

All linear models used in the study were built with the BayesFactor package [54, 55] which implements full Bayesian linear regression. The default Cauchy prior for mean effect size and inverse Wishart prior for effect variance were used. Bayes factors comparing the alternate model containing the variable of interest (usually diagnosis) to the null model without the variable of interest were computed using the default approximation method and log10-transformed. A log10-transformed Bayes factor (logBF) greater than 0.5 was considered significant, a threshold consistent with previous analyses [49]. Posterior distributions for all model parameters were estimated using Gibbs sampling with 10,000 iterations, the median of the distributions was used as the maximum a posteriori (MAP) estimate of each model parameter. When needed, residuals were computed by subtracting the MAP estimated effects of variables from the original data.

Posterior probabilities for individual pairwise contrasts being non-zero were computed by finding the proportion of posterior samples with a sign opposite of the MAP estimate for that parameter. A posterior probability of 0.95 or greater was considered significant. The global FDR for differential expression analyses was computed by taking the average of the posterior probabilities for all genes declared significant for an individual analysis.

#### Differential Expression

Linear models as described above were used to identify differentially expressed (DE) genes in normalized, batch corrected data. The alternate model for each gene included the variable of interest (either diagnosis or APOE allele) along with age and sex as covariates, unless age was confounded, in which case it was removed by residualization before fitting the final model. The null model contained only the covariates. DE genes were defined as genes with a logBF > 0.5 for diagnosis or ApoE allele. Furthermore, in models in which diagnosis had more than 2 values, the posterior probability of the pairwise difference between two diagnoses had to exceed 0.95 to be considered DE. Correlation between differential expression across FTD disorders was computed using biweight midcorrelation of logFC values of each diagnosis vs. control and disease were clustered using average-linked hierarchical clustering.

#### Weighted Gene Co-expression Network Analysis (WGCNA)

We constructed gene co-expression networks using the WGCNA package [15]. The expression data used for each network was normalized and corrected for batch effect but no other covariates were removed. Signed adjacency matrices with a soft power of 12 were computed using biweight midcorrelation [56], converted to topological overlap matrices [52] and clustered using the default average-linked hierarchical clustering. Modules were identified using dynamic tree cutting of the hierarchical clustering tree [57] with cut height of 0.995 and a deepSplit parameter of 2. Module eigengenes were computed from the first principal component of the expression values of the genes in each module, and correlated modules were merged using a dissimilarity threshold of 0.2. We determined eigengene significance using linear models with the same design and significance cutoffs as described above, and removed confounding covariates using residualization on the eigengenes before fitting the final model.

#### Cell type composition

Cell type composition was estimated using support vector regression implemented by CiberSort [24] with the LM22 dataset for immune cells generated for that tool. Cell type compositions were converted from proportions to log odds ratios to give them gaussian distributions and confounding covariates were removed using residualization. The same linear models used for differential expression were used to identify if significant differences in cell type composition could be found between diagnostic groups.

#### Pathway enrichment analysis

Pathway enrichment was analyzed using the GO Biological Process 2017b dataset downloaded from Enrichr [58]. A Bayesian hypergeometric overlap test [59] was used to determine if overlap between a given gene set and all gene sets in the GO Biological Process was significant, with a logBF > 0.5 being defined as significant.

#### Enrichment for genetic risk with MAGMA

We used the MAGMA tool [25] to compute enrichment for genetic risk in the top 300 genes in our co-expression network modules. MAGMA aggregates the genetic association Z-scores for individual SNPs in a given gene into a single gene-level score, and then fits a linear mixed effects model which tests whether membership in a gene set significantly increases the association Z-score while accounting for linkage disequilibrium between genes. The resulting p-value represents the probability of observing the difference in Z-score between the gene set members and non-members under the null hypothesis that the difference is 0. All GWAS studies and annotations used hg19/GRCh37 coordinates. We removed all SNPs within 1 centimorgan of rs7412, the SNP for ApoE2, to eliminate any effect of ApoE. Centimorgan coordinates were taken from a chromosome 19 genetic map generated from 1000 Genomes European samples [60]. Annotation of SNPs to HGNC symbols was done using Ensembl 87, the last release available for hg19, and no window was added up or downstream of genes. Linkage disequilibrium between genes was estimated using 1000 Genomes Phase 3 [60] European samples and synonymous SNPs were dropped. SNP-level association statistics were aggregated using the default mean method.

#### Enrichment for cell type specific expression

We used pSI ([21, 22]) to estimate enrichment of co-expression network modules for cell type specific genes from published data sets in peripheral blood [23] and the CNS [32]. Cell type specific genes were identified by computing the specific index using the default value of 100 permutations considering genes with an expression cutoff of the 5% quantile of all expression values. Genes with a specific index p-value < 0.01 were included in each cell type specific list. In blood, we identified 353 basophil-specific genes, 372 naive B-cell-specific genes, 168 mature B-cell-specific genes, 104 myeloid dendritic cell-specific genes, 44 eosinophil-specific genes, 226 neutrophil-specific genes, 272 megakaryocyte-specific genes, 296 monocyte-specific genes, 66 mature NK-cell-specific genes, 49 memory CD8+ T-cell-specific genes, 187 naive CD8+ T-cell-specific genes, 115 naive CD4+ T-cell-specific genes and 83 memory CD4+ T-cell-specific genes. In the CNS, we identified 433 microglia-specific genes, 343 endothelial cell-specific genes, 400 neuron-specific genes, 295 astrocyte-specific genes, and 196 oligodendrocyte-specific genes. Enrichment was tested using hypergeometric overlap testing as described for pathway enrichment.

#### Enrichment for transcription factor binding

We used an approach very similar to that used by MAGMA [25] adapted for working with ChIP-Seq data. We downloaded bigWig files in GRCh38 coordinates for the signal log p-values computed using both technical replicates for Spi1/PU.1 (HL-60, ENCFF935IFD), RELA (K562, ENCC460WXB), CEBPB (K562, ENCFF999NRK), JUN (K562, ENCC243LNJ), RUNX1/AML1 (K562, ENCFF825MDH) and NFYA (K562, ENCFF180VKB) from ENCODE (https://www.encodeproject.org). Gene annotations were taken from the latest version of Ensembl for GRCh38 (Ensembl 93), and a 1.5 kb window was added upstream of all transcription start sites to cover promoter regions. Log p-values of all peaks in each gene window were averaged for all genes expressed in co-expression networks. Enrichment testing of each module was performed using a linear model with a dummy variable for the top 500 genes in each module, or all genes if the module had less than 500 genes.

### Data and software availability

All gene expression microarray data is available for download from the Gene Expression Omnibus (https://www.ncbi.nlm.nih.gov/geo/query/acc.cgi?acc=GSEXXXXX). All other data used in the study was previously published and is available from the linked websites. All R packages used in statistical analysis are available in CRAN or BioConductor, and all other software used is freely available from its linked website.

---

**Table S1. Tables of differential expression (DE) statistics for each gene in the analysis for all AD, MCI, and control subjects, sex-specific analyses of AD, MCI and control subjects, and FTD and control subjects. Related to Figure 1.** The first two columns show the HGNC symbol and gene description and the third column is the logBF for the model comparison. Subsequent columns show logFC and posterior probabilities for each pairwise comparison.

**Table S2. Tables with network information for eighted gene co-expression network analysis (WGCNA) for the network of all AD, MCI, and control subjects, sex-specific networks of AD, MCI and control subjects, and the network of FTD and control subjects. Related to Figure 2 and Figure 3.** The first three columns show the HGNC symbol, description and module assignment for each gene. The next 5 columns show connectivity measures: total connectivity (kTotal), within module connectivity (kWithin), outside module connectivity (kOut), difference between within and outside module connectivity (kDiff), and scaled within module connectivity (kscaled). The subsequent columns show the module membership of each gene in each module.

**Table S3. Cell type enrichment analysis for all blood cell types in WGCNA modules. Related to Figure 4 and Figure 6.** Tables of logBF of overlap between the top 300 genes in each module in the AD and FTD networks and cell types specific genes in blood and CNS.

**Table S4. Disease risk enrichment analysis of WGCNA modules for additional GWAS studies. Related to Figure 5.** Tables of FDR-adjusted p-values of significance of enrichment of the top 300 genes in each module in the AD and FTD networks for genetic risk in ALS, diabetes, Huntington’s Disease, major depression and schizophrenia.

**Table S5. Binding scores for PU.1/SPI1, RELA, CEBPB, JUN, RUNX1, and NFYA for all genes. Related to Figure 7.** The first four columns show the gene name, chromosome, start, and end under hg38 coordinates. The remaining columns are scores for each transcription factor.

**Figure S1.**
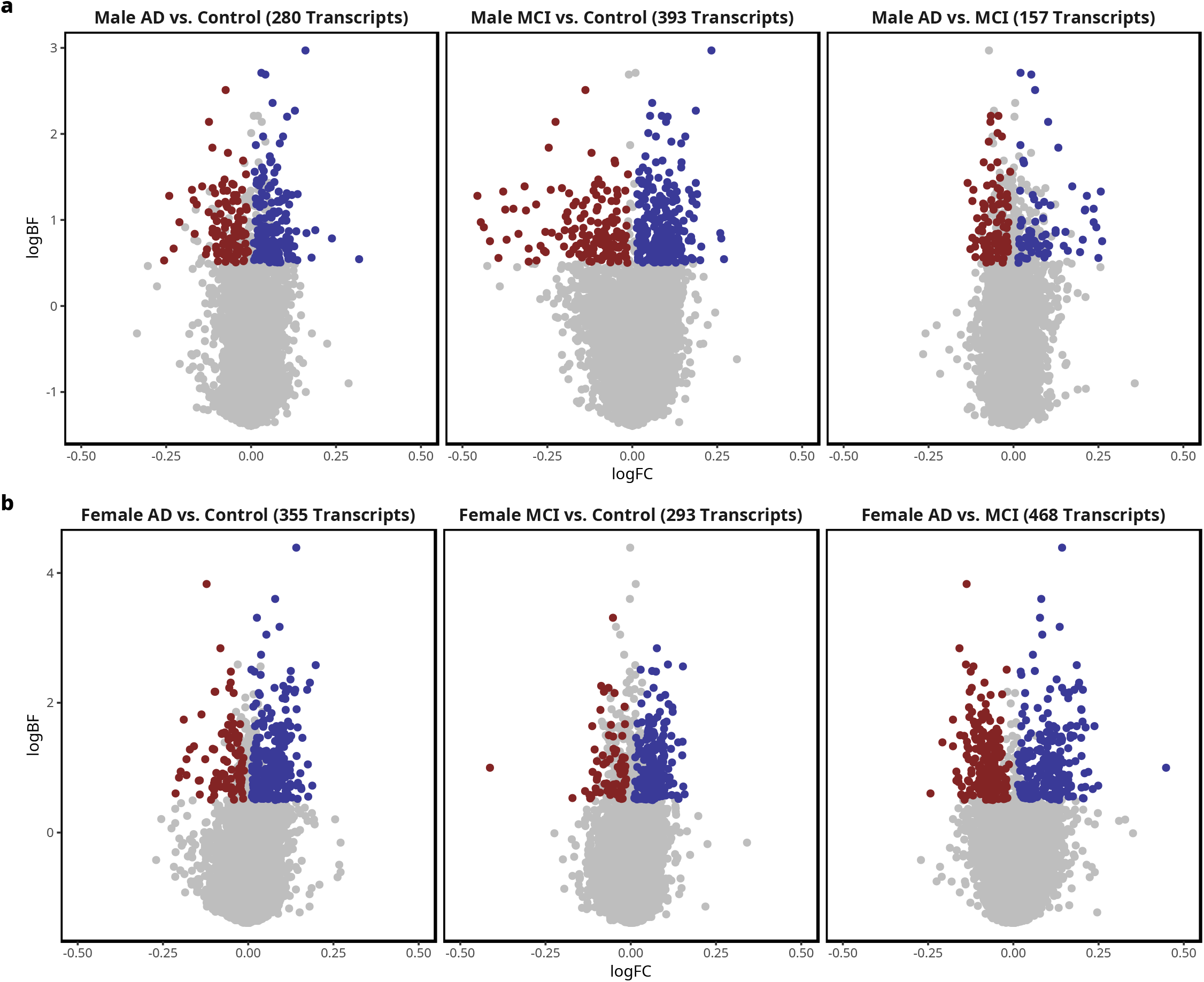
Differential expression analysis of sex-specific AD, MCI, and control cohorts. Related to Figure 1. **a,b** Volcano plots of the log fold change (logFC) in gene expression on the x-axis versus the log10 Bayes Factor (logBF) on the y-axis, with downregulated genes in blue and upregulated genes in red. Only genes with a significant pairwise probability greater than 0.95 were colored. The contrast and number of DE genes are shown in the plot titles.

**Figure S2.**
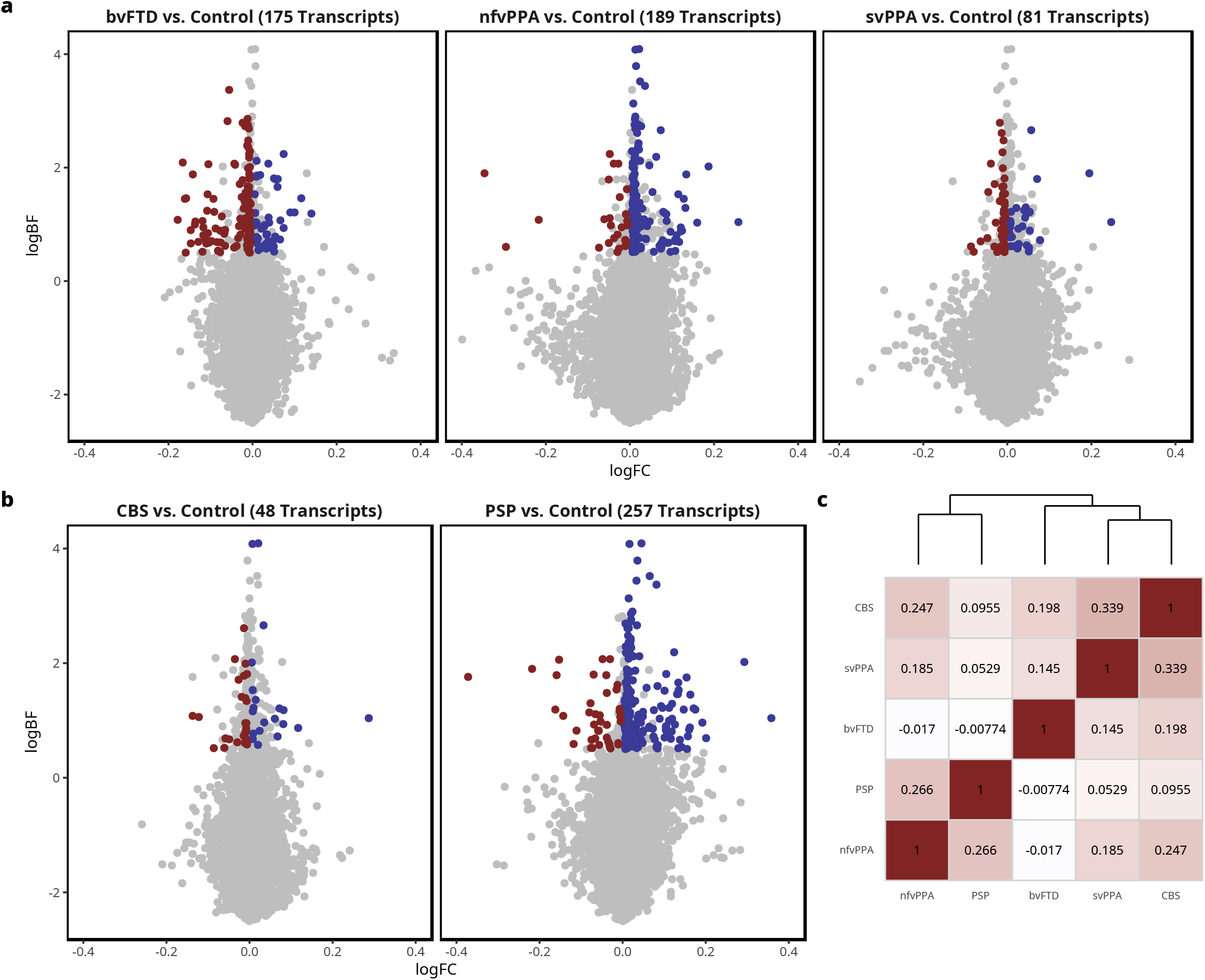
Differential expression analysis of FTD disorders vs. controls. Related to Figure 1. **a,b** Volcano plots of the log fold change (logFC) in gene expression on the x-axis versus the log10 Bayes Factor (logBF) on the y-axis, with downregulated genes in blue and upregulated genes in red. Only genes with a significant pairwise probability greater than 0.95 were colored. The contrast and number of DE genes are shown in the plot titles. **c** Correlation plot of the pairwise correlation of logFC values of FTD disorders vs. control, with the dendrogram on top showing hierarchical clustering of disorders.

**Figure S3.**
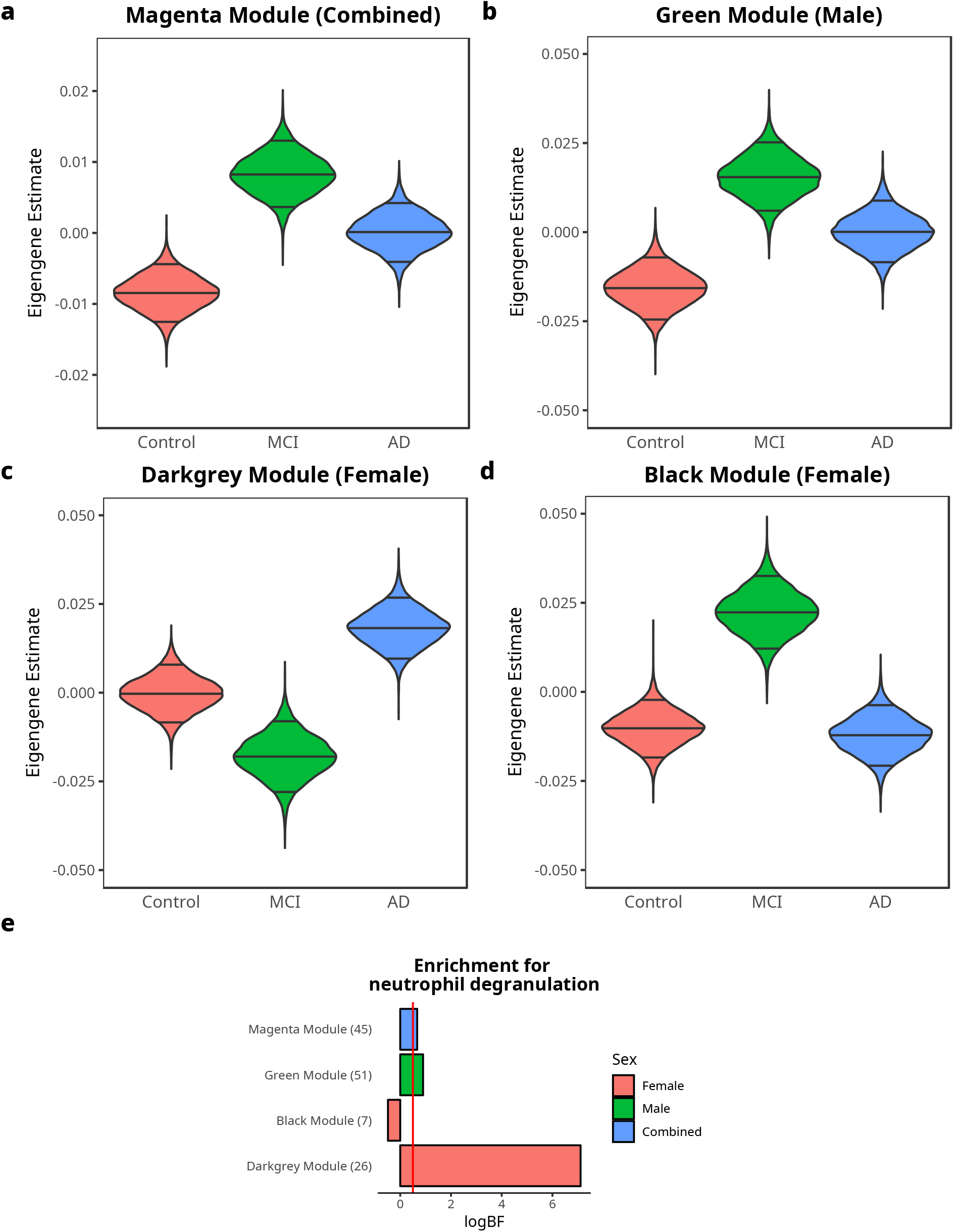
Additional WGCNA modules significantly associated with disease. Related to Figure 2. **a-d** Violin plots of posterior estimate of mean eigengene values for each diagnosis, with the median and 5% and 95% quantiles indicated by lines. **e** Bar plot of enrichment of genes in each module for neutrophil degranulation (GO:0043312) with the logBF on the x-axis.

**Figure S4.**
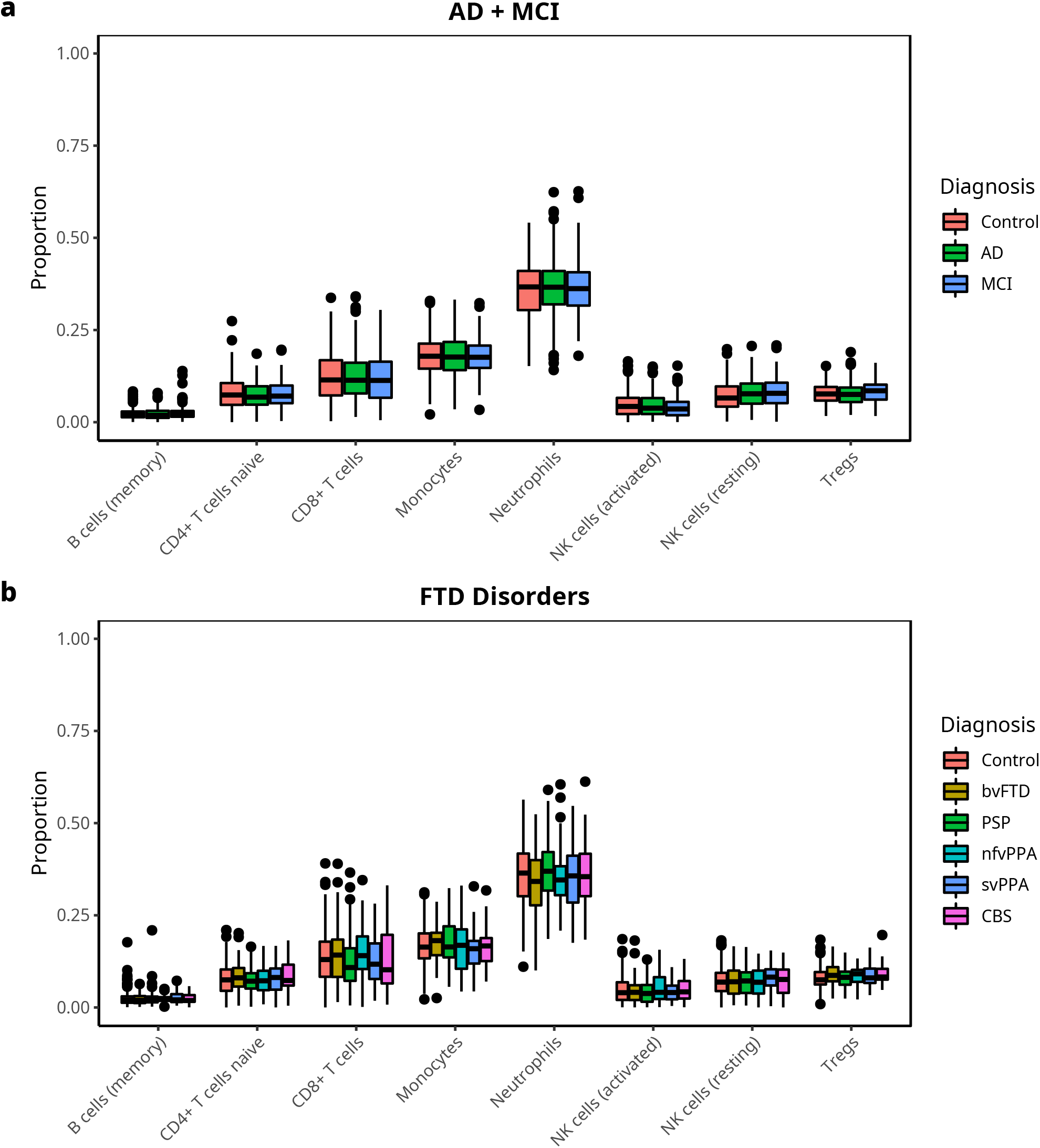
Cell type composition analysis reveals no significant differences in cell between disease and control groups. Related to Cell Type Composition section. **a,b** Box plots of blood cell type composition estimated from gene expression in AD, MCI and control (**a**) and FTD disorders (**b**).

**Figure S5.**
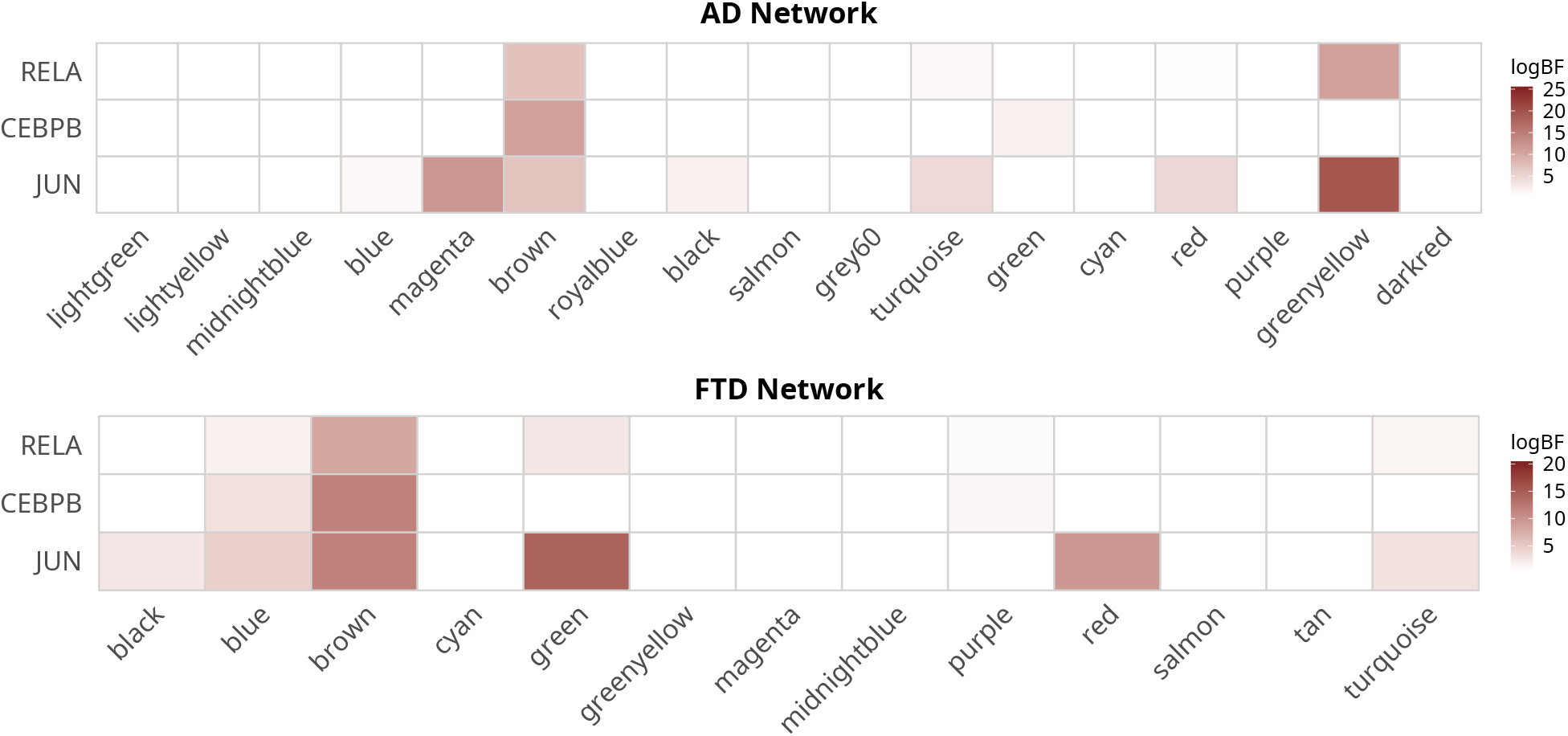
Binding enrichment analysis of WGCNA modules reveals additional specific enrichment in disease-associated modules. Related to Figure 7. Heatmap of enrichment for binding of transcription factors for the top 500 genes in modules in the AD and FTD networks.

